# Early Antibody-Mediated Immunity to Tuberculosis in Mice Requires NLRP3

**DOI:** 10.1101/2025.07.18.665318

**Authors:** Rania Bouzeyen, Nyaradzai Sithole, Avia Watson, Lilach Abramovitz, Katya Rakayev, Adam Fillion, Tessa Dickson, Paul A. MacAry, Patricia A. Darrah, Robert A. Seder, Natalia T Freund, Babak Javid

## Abstract

While antibodies have emerged as potential mediators of protective immunity against *Mycobacterium tuberculosis* (Mtb), their mechanisms of action remain incompletely understood. Here, we demonstrate that immune complexes of Mtb and monoclonal antibodies targeting the Mtb phosphate transporter subunit PstS1 robustly activate the NLRP3 inflammasome in human and murine macrophages, leading to enhanced interleukin-1β secretion. Surprisingly, antibody-mediated inflammasome activation occurred independently of cell-surface Fcγ receptors, as confirmed using Fc-domain glycosylation mutant mAbs and macrophages from Fcγ receptor-deficient mice. Crucially, NLRP3 is indispensable for early antibody-mediated protection in vivo, as both pharmacological inhibition, and genetic deletion of NLRP3 completely abolished protective effects of PstS1-specific antibodies in Mtb-infected mice. This mechanism extends beyond monoclonal antibodies, as polyclonal sera from intravenously BCG-immunized rhesus macaques also required NLRP3 for protective efficacy. Our findings reveal a previously unrecognized mechanism by which Mtb-specific antibodies enhance host defense through inflammasome activation, potentially informing novel approaches for tuberculosis vaccine development.

## Main text

Tuberculosis (TB), caused by *Mycobacterium tuberculosis* (Mtb), remains the world’s deadliest infectious disease, with ∼10 million cases of active TB per year, resulting in ∼1.3 million deaths. The only licensed TB vaccine, Bacillus Calmette–Guérin (BCG) fails to protect against pulmonary TB, that drives continued transmission worldwide^1^. While cell-mediated immunity is established as crucial for TB control, mounting evidence supports a protective role for antibody-mediated immunity^2, 3, 4, 5, 6, 7, 8^. Post-hoc analysis of the M72/AS01_E_ subunit vaccine, which showed ∼50% efficacy, implicated Mtb-specific antibody responses as a potential correlate of protection^9^. However, the mechanisms by which Mtb-specific antibodies confer protection are not fully understood^10^.

We previously isolated and characterized monoclonal antibodies (mAbs) to the Mtb phosphate transporter subunit PstS1 from a patient with active TB. Two PstS1-specific mAbs, p4-36 and p4-163, that bind distinct epitopes, restricted Mtb growth in experimental infection models including early murine infection^7^. Here, we used these PstS1-specific mAbs to investigate mechanisms by which Mtb-specific antibodies mediate protective immune responses in macrophage infection and in mice.

### Mtb-specific mAbs complexed to Mtb stimulate inflammasome-dependent IL-1β secretion

To investigate mechanisms underlying antibody-mediated protection against *Mycobacterium tuberculosis* (Mtb), we focused on inflammasome activation, a critical pathway in innate immunity. Inflammasomes are multiprotein complexes^11^ that trigger the secretion of pro-inflammatory cytokines IL-1β and IL-18. While Mtb can stimulate the NLRP3 inflammasome ^12, 13, 14^, the pathogen has evolved numerous evasion mechanisms^15, 16, 17, 18, 19, 20, 21, 22^, leaving the precise role of inflammasomes in early tuberculosis immunity unresolved^13, 23^.

Previous work had suggested that polyclonal antibodies from Mtb-exposed individuals might induce ASC (Apoptosis-associated speck-like protein containing a Caspase activation and recruitment domain) speck formation^6^, a hallmark of inflammasome activation^24^. Building on this observation, we found that PstS1-specific monoclonal antibodies (mAbs) p4-36 and p4-163, when complexed with Mtb (Mtb/Ab), significantly enhanced ASC speck formation in differentiated THP-1 cells compared with an irrelevant IgG1 isotype control (**Fig. 1a, b**). This effect translated to robust IL-1β secretion – a canonical product of inflammasome activation – in differentiated THP-1 cells, primary human monocyte-derived macrophages (HMDM), and murine bone marrow-derived macrophages (BMDMs) (**Fig. 1c-e**).

**Figure 1.**
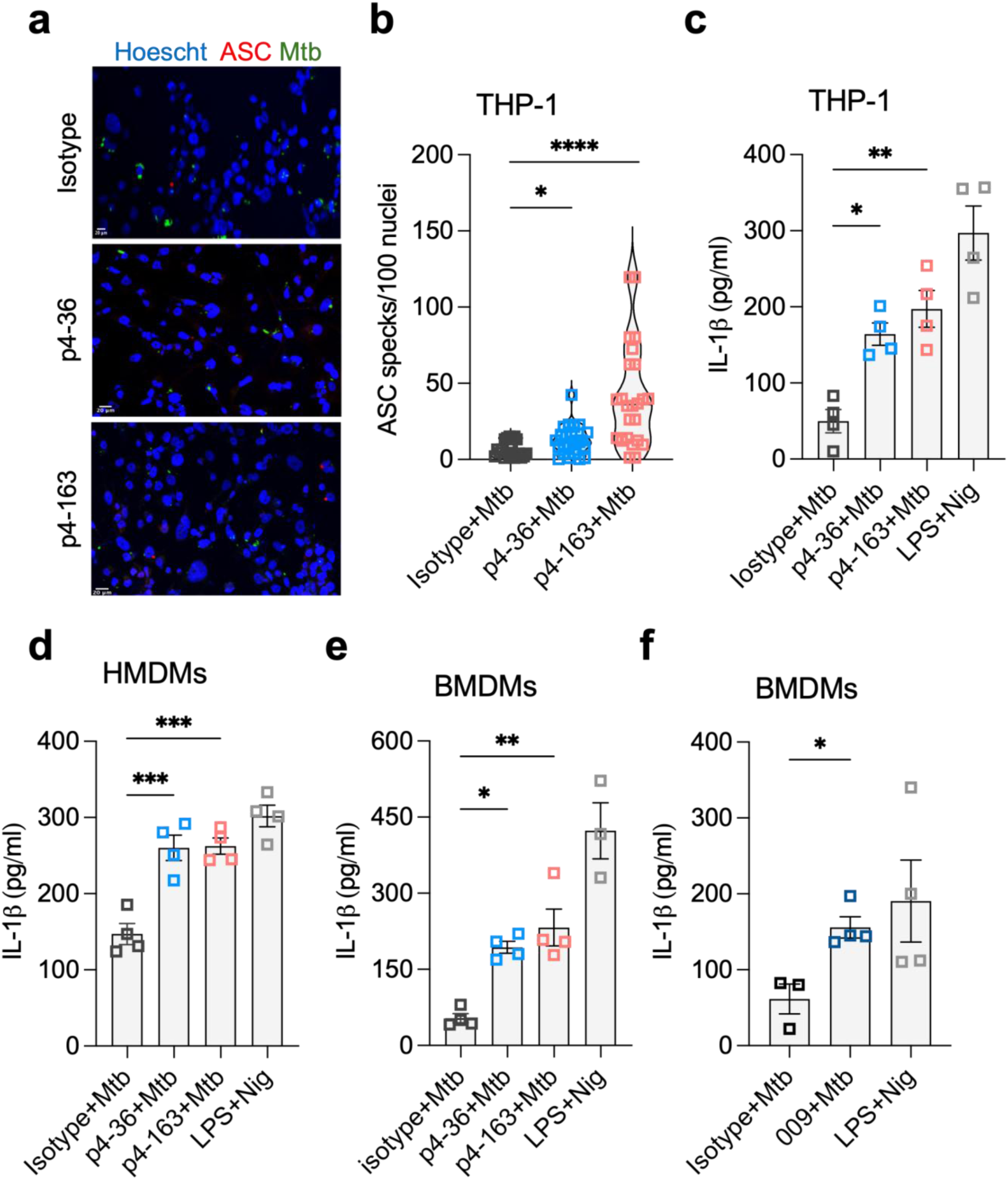
Mtb-specific mAbs complexed to Mtb stimulate inflammasome-mediated IL-1β secretion. (a) ASC speck formation visualized by confocal microscopy. PMA-differentiated THP-1 macrophage-like cells were incubated with Mtb-zsGreen/Ab (p4-36, p4-163 or IgG1 isotype control – see Methods) immune complexes at an MOI of 5:1. Cells were stained, fixed and imaged after 24hrs. ASC specks, red, were quantified/ 100 nuclei (stained with Hoescht). Representative high-power field images from the three experimental conditions shown. Scale bar 20 µm. (b) Quantification of the analysis in (a). Each dot represents data from an independent experiment. PMA-differentiated THP-1 cells (c), primary human monocyte-derived macrophages (d) and bone-marrow derived macrophages from C57BL/6 mice (e, f) were infected with Mtb/Ab immune complexes (mAb concentration 25µg/mL) at an MOI of 1 (THP-1) and 5 (BMDM) respectively. LPS + Nigericin (LPS+Nig) used as a positive control from inflammasome activation. Supernatants from culture medium were collected at 24hrs post-infection and assayed for IL-1β by ELISA. Panels c-f data representative of 5 independent experiments and shown as mean +/- SEM. Each dot represents data from a biological replicate. *p<0.05, **p<0.01, ***p<0.001, ****p<0.0001 by one-way ANOVA and Dunnett’s post-hoc correction for multiple comparisons.

Recombinant PstS1 complexed with PstS1-specific mAbs failed to stimulate IL-1β secretion (**Extended Data Fig. 1**), indicating that inflammasome activation requires antibody engagement with intact Mtb rather than soluble antigen alone. Importantly, this phenomenon was not restricted to PstS1-specific antibodies, as an LpqH-specific mAb, (009), we recently isolated^25^ from an individual with occupational TB exposure^4^, similarly enhanced IL-1β secretion from infected BMDMs (**Fig. 1f**), confirming that inflammasome activation by Mtb/Ab was generalizable across distinct antigen targets.

### Mtb/antibody complexes activate the canonical NLRP3 inflammasome pathway

Having established that Mtb/Ab complexes trigger IL-1β secretion, we next sought to identify which specific inflammasome pathway mediates this response. To characterize the specific inflammasome components required for antibody-enhanced immunity we measured Caspase-1 activation, which is required for canonical inflammasome activation. Mtb/PstS1-specific mAbs significantly increased active Caspase-1 compared with IgG1 isotype control, as measured by the fluorescent inhibitor FLICA (**Fig 2a,b**). Both pharmacological inhibition using the Caspase-1 inhibitor VX765 and genetic deletion of Caspase-1 in THP-1 cells abolished antibody-mediated IL-1β secretion (**Fig. 2c,d**), confirming canonical inflammasome pathway involvement.

**Figure 2.**
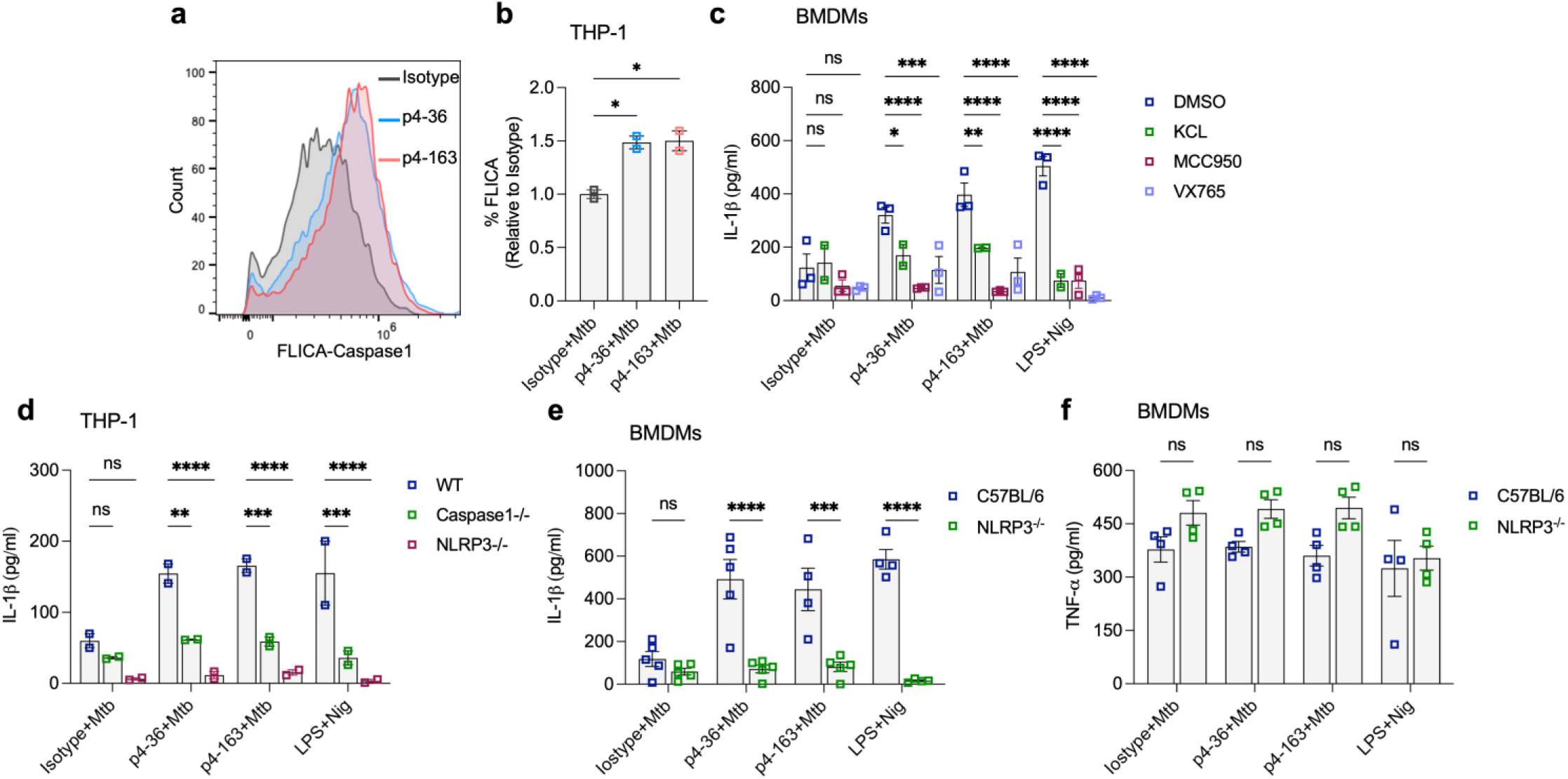
Mtb/PstS1 mAb complexes stimulate IL-1β via NLRP3 inflammasome activation. (a) PMA-differentiated THP-1 cells were infected with Mtb/Ab (mAb concentration 25µg/mL) at MOI of 1. After 24hrs, cells were stained with FAM-YVAD-FMK probe (FLICA) and analyzed by flow cytometry. (b) The proportion of FLICA+ cells from two independent experiments from each experimental group shown as mean +/- SEM. IL-1β in culture supernatant at 24hrs post-infection from BMDM – derived either from wild-type C57BL/6 or *Nlrp3^-/-^* mice as indicated (c, e) and PMA-differentiated THP-1 cells – wild-type, *Caspase-1*-deleted or *Nlrp3*-deleted as indicated (d). Cells were infected with Mtb/Ab immune complexes, MOI 5 (BMDMs) or 1 (THP-1). Cells were treated with VX765 (50 µM), MCC950 (10 µM) or with KCl (5 mM) in the culture medium, or vehicle control (c). (f) TNFα in culture supernatant at 24hrs post-infection as measured by ELISA in same experiment as (e). Data shown as mean +/- SEM. Each dot represents data from a biological replicate. ns p>0.05, *p<0.05, **p<0.01, ***p<0.001, ****p<0.0001 by one-way ANOVA and Dunnett’s post-hoc correction for multiple comparisons.

Having established Caspase-1 dependence, we next identified the upstream sensor responsible for inflammasome assembly. Since NLRP3 is a key inflammasome sensor stimulated by Mtb infection^12, 14, 18, 26^, we tested its requirement in Mtb/Ab-mediated IL-1β secretion. Both the NLRP3-specific inhibitor MCC950^27^ and potassium chloride in the culture medium, which blocks NLRP3 activation, abrogated IL-1β secretion (**Fig. 2c**). As further confirmation, infection of NLRP3-deleted THP-1 cells (**Fig. 2d**) and BMDMs (**Fig. 2e**) completely prevented IL-1β secretion, while preserving secretion of the inflammasome-independent cytokine TNFα (**Fig. 2f**), highlighting the specificity of the effect for inflammasome-dependent responses.

Together, these data demonstrate that under infection conditions in which Mtb alone minimally induced inflammasome assembly, Mtb-specific mAbs complexed to Mtb potently stimulate NLRP3-dependent inflammasome activation and IL-1β secretion.

### Antibody-mediated inflammasome activation occurs independently of cell-surface FcγRs

Having demonstrated NLRP3-dependent inflammasome activation by Mtb/Ab complexes, we investigated the upstream receptor mechanisms. Since antibody effector functions classically depend on Fc-FcγR interactions^28^, we hypothesized that FcγRs mediate this inflammasome activation. Antibody engineering enables focusing of Fc-effector functions via mutation of the Fc antibody domain, to increase/decrease engagement of specific receptors^29, 30, 31^. Surprisingly, Mtb/Ab immune complexes with an IgG1-N297A mAb variant, deficient in FcγR binding due to abolished Fc glycosylation, induced IL-1β secretion to the same extent as wild-type IgG1 (**Fig. 3a, b**). This indicated that classical cell-surface FcγR engagement is dispensable for this antibody-mediated inflammasome activation.

**Figure 3.**
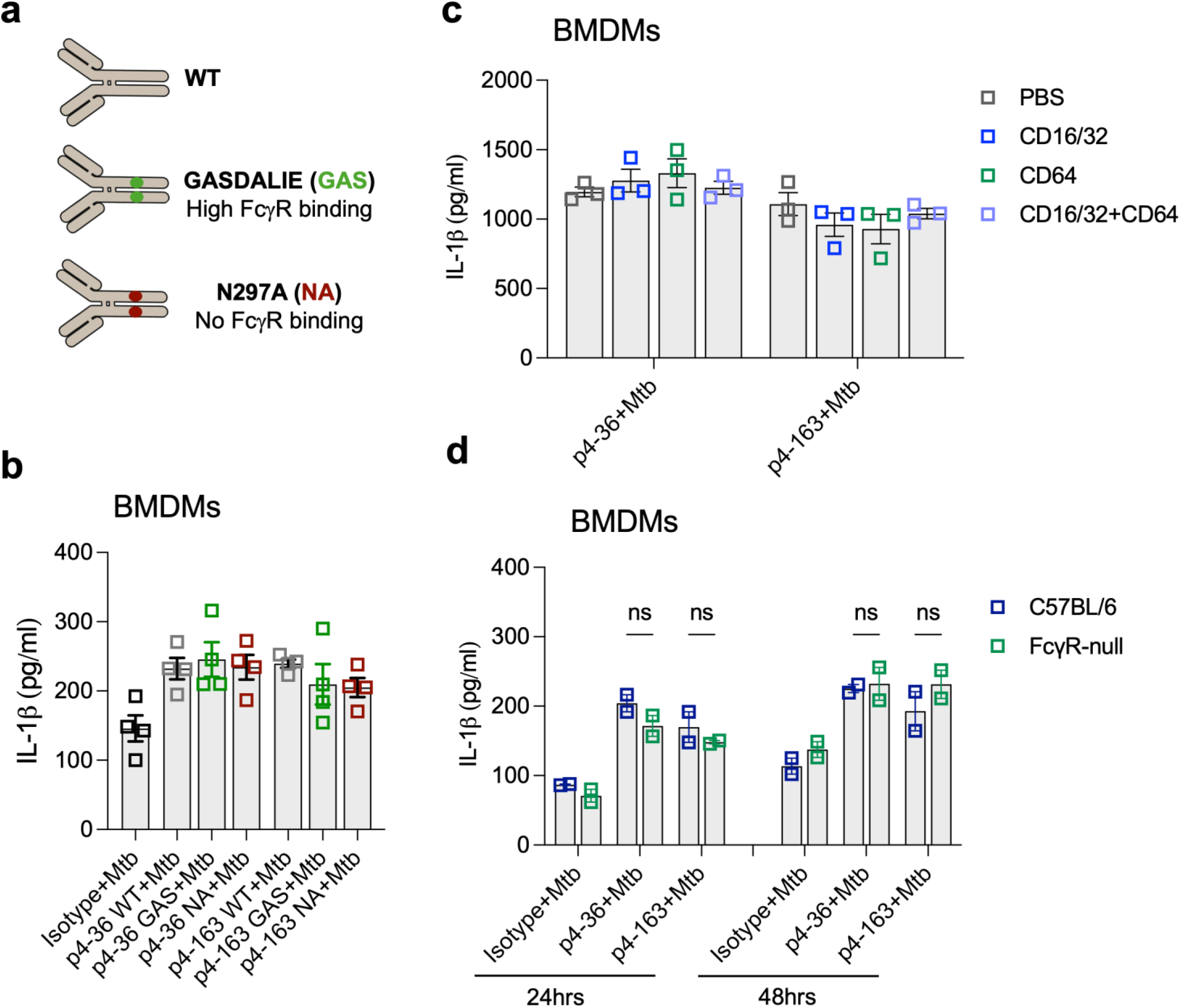
Antibody-mediated inflammasome activation occurs independently of cell-surface FcγRs. (a) Cartoon of engineered human IgG1 mAbs. Fc mutants: GASDALIE (GAS) – G236A/S239D/A330L/I332E IgG1 has increased engagement of activatory FcγRs and N297A (NA) eliminates the sole N-glycosylation site of the Fc and abolishes FcγR binding. (b-d) IL-1β as measured by ELISA at 24hrs from culture supernatants of BMDMs infected with Mtb/Ab immune complexes (MOI 5). (c) Prior to infection with Mtb/Ab immune complexes, macrophages were treated with anti-CD16 (CD16), anti-CD32 (CD32) and CD64 (CD64) at 5µg/mL concentration either solely or in combination. (d) BMDMs from C57BL/6 or FcγR-null mice were infected with Mtb/Ab as before except that IL-1β was assayed at 48hrs as well as 24hrs post-infection. Each dot represents data from a biological replicate. Data shown as mean +/- SEM and representative of two independent experiments. ns p>0.05, **p<0.01, ***p<0.001, ****p<0.0001 by two-way ANOVA and Sidak’s post-hoc correction for multiple comparisons.

Blocking the FcγRs CD16, CD32 and CD64 individually or together with specific antibodies also failed to inhibit IL-1β secretion (**Fig. 3c**). Furthermore, BMDMs from mice lacking the FcγR common chain, “FcγR null” mice^32^, were as competent as wild-type BMDMs for Mtb/Ab-mediated IL-1β secretion (**Fig. 3d**). These multiple orthogonal approaches confirm that FcγRs are not required for Mtb/Ab-mediated IL-1β secretion.

### NLRP3 is required for early antibody-mediated protection to Mtb in vivo

Our in vitro findings suggested that NLRP3 inflammasome activation underlies antibody-mediated immunity against Mtb. To test the physiological relevance of these findings and to determine whether this mechanism operates in vivo, we hypothesized that NLRP3 would be required for antibody-mediated protection in Mtb-infected mice. We expanded our earlier studies of PstS1-specific mAb-mediated immunity in mice^7^ with dose-ranging studies. Administering 0.05-0.5 mg anti-PstS1 mAb/mouse intra-peritoneally, one day prior to aerosol infection with Mtb (**Fig. 4a**), we identified a dose of 0.1mg/ mouse offered optimal protection for both PstS1-specific antibodies (**Fig. 4b**).

**Figure 4.**
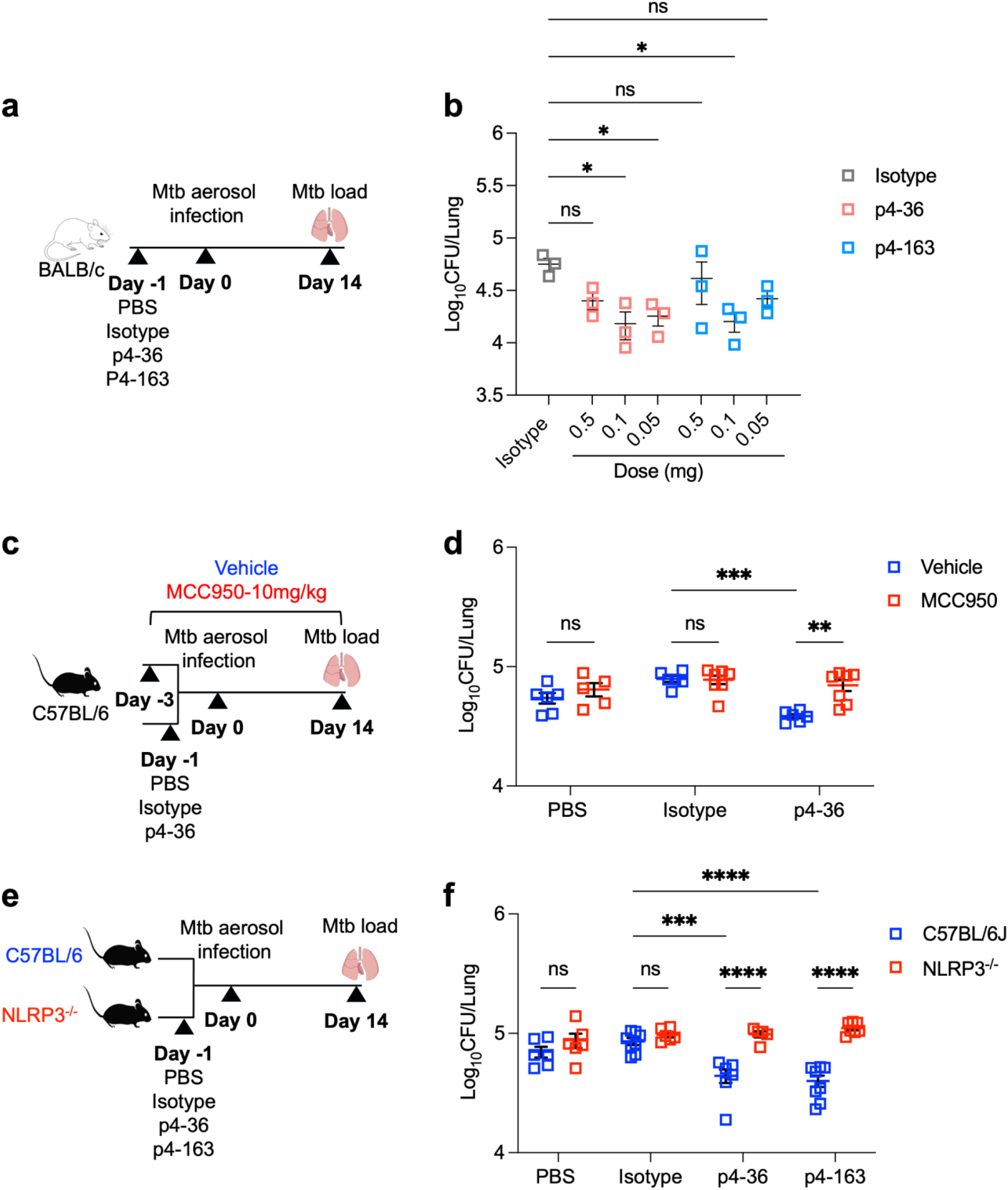
NLRP3 is required for early antibody-mediated immunity to Mtb in vivo. (a) Cartoon schematic of experimental design in dose-ranging experiment. Balb/C mce were injected intraperitoneally with indicated dose of mAb per animal (p4-36, p4-163 or isotype IgG1) 24hrs prior to aerosol infection with Mtb-HN878 – delivered dose 20-30 CFU/animal. Lung homogenates of animals euthanized at day 14 post-infection were plated for CFU (b). (c) Cartoon schematic of experimental design. C57BL/6 were treated with daily injections of MCC950 (10mg/kg) or vehicle from 3 days prior to infection. One day prior to infection with Mtb-HN878 (10-20 CFU delivered dose/animal), mice were injected with 0.1mg mAb/mouse as before. CFU from lung homogenates at day 14 enumerated as before (d). (e) Cartoon schematic of experimental design. C57BL/6 or *Nlrp3^-/-^* mice were injected with 0.1mg of mAb/mouse one day prior to aerosol infection with Mtb-HN878 (delivered dose 15-20 CFU/animal) as before. CFU from lung homogenates at day 14 enumerated as before. Data are representative of two separate experiments and each dot represents CFU from one animal, with mean as horizontal bar. ns p>0.05, *p<0.05, **p<0.01, ***p<0.001, ****p<0.0001 by one-way ANOVA and Dunnett’s post-hoc correction for multiple comparisons.

We then tested whether NLRP3 is required for antibody-mediated protection in Mtb-infected mice. Pretreatment with the NLRP3 inhibitor MCC950 had no effect on bacterial burden in mock (saline) or isotype control-treated mice, consistent with previous studies showing no role for NLRP3 in TB immunity in naïve animals^6^. Strikingly, however, MCC950 completely abrogated the CFU reduction observed at day 14 post-infection in p4-36-treated mice (**Fig. 4c,d**). To confirm this finding was not due to off-target effects, we tested both p4-36 and p4-163 antibodies in *Nlrp3*-knockout mice. Again, while NLRP3 deletion did not increase susceptibility to Mtb in naïve mice, it completely abolished the protective effect of Mtb-specific antibodies (**Fig 4e,f**).

Cytokines and chemokine analysis of lung homogenates from wild-type and *Nlrp3*^-/-^ Mtb-infected mice at day 14 revealed that several TB immunity-associated cytokines, such as TNFα, did not vary by condition, whereas others, such as IL-6 varied between wild-type and *Nlrp3*^-/-^ mice, but not in a PstS1 mAb-specific manner (**Extended Data Fig. 2**). Bulk lung IL-1β or IL-18 levels did not significantly differ by NLRP3 status at this time point (**Extended Data Fig. 2**), possibly reflecting contributions from non-inflammasome sources^23^, or localized effects not captured in whole lung homogenates. These observations suggest that the protective effect of antibody-NLRP3 engagement may manifest through early, localized, or dynamic cellular responses that are not fully reflected in bulk cytokine levels at day 14, by which time a significant reduction in bacterial burden is already established.

To further extend our studies beyond monoclonal antibodies, we tested sera from rhesus macaques that had been immunized with BCG either via the intradermal (I.D.) or intravenous (I.V.) routes (**Extended Data Table 1**). Recent work has shown that intravenous, but not intradermal BCG immunization provides almost complete protection to Rhesus macaques against Mtb challenge^33^. Post-hoc analyses to identify potential mediators of this IV-BCG-mediated protection identified both antibody and cell-mediated responses as biomarkers^34, 35^, although the causality of antibody-mediated responses in protection were less clear ^34^. Total plasma immunoglobulin was similar in all animals (**Extended Data Fig. 3a, b**). Plasma from IV-immunized animals had greater binding to BCG and heat-killed Mtb-HN878, but not to *E. coli* (**Extended Data Fig. 3c-e**). When complexed to Mtb and used to infect macrophages, both sera from intradermally- and intravenously-immunized animals were able to stimulate IL-1β secretion, with sera from IV-immunized animals significantly more potent than sera from ID-immunized macaques (**Fig. 5a**). Crucially, passive transfer of sera from macaques protectively vaccinated with intravenous BCG also conferred modest early protection against Mtb challenge in wild-type mice (**Extended Data Fig. 3f, g**). This protection was entirely abrogated in *Nlrp3*^-/-^ mice, demonstrating that NLRP3 is critical for the efficacy of these vaccine-induced polyclonal antibodies (**Fig. 5b,c**).

**Figure 5.**
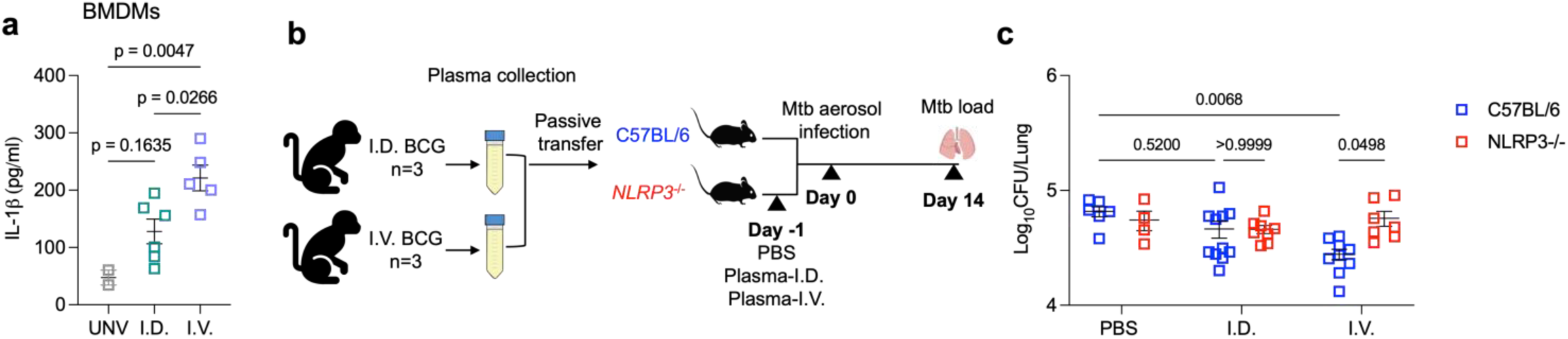
Immunity to tuberculosis from passive transfer of macaque plasma from intravenously-BCG-immunized animals depends on NLRP3. (a) BMDM were infected (MOI 5) with Mtb/Ab immune complex with plasma derived from unvaccinated (UNV), intradermally- (ID) or intravenously- (IV) BCG-immunized Rhesus macaques. 24hrs following infection, IL-1β from supernatants was measured. (b) Cartoon schematic of macaque plasma passive transfer experiment. (c) Plasma (0.5 mL) from ID- (n=3) and IV- (n=3) immunized animals was injected into wild-type C57/BL6 or *Nlrp3*^-/-^ mice 1 day prior to aerosol challenge with Mtb-HN878 (∼10 cfu/animal). Donor plasma from each macaque was injected into 2-3 recipient mice. Bacterial cfu from lung homogenates from euthanized mice at d14 are shown. Each dot represents data from a single mouse and bar represents mean. p-values from one-way ANOVA and post-hoc correction shown.

## Conclusions

Our study identifies a novel FcγR-independent mechanism by which Mtb-specific antibodies promote host defense, demonstrating that antibody-Mtb complexes directly trigger the NLRP3 inflammasome, leading to IL-1β secretion and enhanced bacterial control. This finding is particularly noteworthy as most antibody effector functions against intracellular pathogens are attributed to FcγR engagement. The ability of pathogen-specific antibodies to operate via an alternative, inflammasome-dependent pathway suggests new mechanisms by which antibodies contribute to host defense. A recent study identified SARS-CoV-2 immune complexes are capable of stimulating inflammasome signaling via FcγRs ^36^. Although the physiological relevance of that finding is less clear, these findings collectively suggest that Ab/pathogen immune complexes may stimulate inflammasome assembly via non-redundant pathways. The physiological relevance of the pathway we identified was confirmed in vivo, where NLRP3 was indispensable for the early protective effects mediated by both monoclonal PstS1-specific antibodies and, significantly, by polyclonal sera from IV BCG-vaccinated Rhesus macaques.

Our discovery of an FcγR-independent pathway for early antibody-mediated protection to tuberculosis does not preclude a role for FcγRs in immunity. Work by Chan and colleagues revealed a role for FcγRs in immunity to Mtb in chronic (>30 days post-infection) but not acute (≤ 20 days post-infection) in mice^37^ and a more recent study^38^ of an engineered LAM-specific mAb showed that N297A-engineered antibodies lost protection in passive transfer. While the precise host receptor that senses Mtb/Ab to initiate this FcγR-independent NLRP3 activation remains to be identified, our data point towards a non-canonical sensing mechanism, potentially involving intracellular receptors like TRIM21^39^. Elucidating this initial sensing step is a key future direction. Mtb is adept at subverting inflammasome activation. Our discovery that Mtb-specific antibodies can overcome this subversion to harness NLRP3 highlights a previously unappreciated layer of antibody-mediated control in TB. These insights suggest that the capacity to elicit antibodies that effectively engage this protective inflammasome pathway could be a critical, and perhaps overlooked, feature of novel TB vaccines. Strategies aimed at inducing such antibody qualities may therefore hold promise for improving vaccine efficacy against this globally important pathogen.

## Methods

### Cell-lines and primary cells

THP-1 cells were from ATCC. Caspase-1 KO cells ^40^ were a gift from Daniel Bachovchin (Memorial Sloan Kettering) while NLRP3-deleted THP-1 cells were purchased from Invivogen (#thp-konlrp3z). THP-1 cells were maintained in RPMI +10% FBS and 1x non-essential amino acids (RPMI-10). For differentiation, cells were seeded for assays in the presence of PMA (10ng/ml) for 48hrs. Cells were washed and rested for 24hrs prior to further experiments. Primary human monocytes were differentiated into macrophages as before^41^ and were a gift from Sara Suliman (UCSF) and were purchased from StemCell.

Bone Marrow derived macrophages (BMDMs) were isolated from femurs of euthanized C57BL/6, *Nlrp3^-/-^*or FcγR-null mice and were cultured and differentiated for six days in DMEM 10% FBS (DMEM-10) and 1x non-essential amino acids in the presence of 20ng/ml murine M-CSF.

### Antibodies and reagents

Monoclonal antibodies targeting PstS1were expressed as human IgG1 and generated as previously described^7^. N297A- and GASDALIE-engineered mAbs were made as previously described^7^. Monoclonal antibody 009 targeting LpqH was expressed as human IgG1 and generated as previously described^25^. Human IgG1 isotype control (BE0297) was purchased from BIOXcell.

### *Mycobacterium tuberculosis* culture

In keeping with prior studies^4, 42^, Mtb was grown in detergent-free conditions for macrophage infection. Growth in the absence of detergent increased mAb binding to Mtb compared with Mtb grown in the presence of detergent (Tween) – **Extended Data Fig. 4**.

For in vitro experiments, Mtb was grown on Middlebrook 7H11 agar, passed through a cell strainer, aliquoted and stored at −80 C in 25% glycerol. Prior to macrophage infection, Mtb stocks were thawed, washed twice with 1X PBS and then passed 10 times through a 29-gauge syringe to remove clumps. For mouse infection, Mtb stocks were grown in Middlebrook 7H9 with ADC supplement and 0.05% Tween-80 at 37 C with constant agitation to an OD=0.8-1. Cultures were then washed, aliquoted and stored at −80 C in 25% glycerol until use.

### Infection and cell treatment

All experiments with Mtb were conducted in the UCSF BSL-3/ A-BSL-3 facilities following institutionally approved biosafety protocols. Single-cell suspension of Mtb-HN878 or Mtb-HN878-zsGreen (for microscopy experiments) grown detergent free were used to infect macrophages at a multiplicity of infection (MOI) of 1:1 Mtb: macrophages for human macrophages and 5:1 for BMDMs. Mtb/Ab immune complexes were pre-formed by incubating bacteria with mAb (25 µg/mL) or isotype control for 1-hour at 37C prior to infection.

Macrophages were infected for 4hrs in serum-free medium prior to washing and replacement with medium containing amikacin for 1hr (250µg/mL; Sigma) to kill extracellular bacteria. Then, cells were washed and overlaid with RPM10 or DMEM10 depending on cell-type. As a positive control for inflammasome induction, cells were primed with 250ug/ml of pure LPS (*E. col*i 0111:B4, Sigma) and 5uM Nigericin (Roche).

### ELISAs

ELISA kits to measure human IL-1β (DY201 and DY401 respectively) were purchased from R&D systems. Mouse ELISA kits for IL-18, IL-10, TNFα, IL-12p70, IL-6, CXCL1, CxCL2 were purchased from Invitrogen. ELISA kit to measure mouse IFN-β was purchased from Biolegend. All assays were performed according to the manufacturer’s instructions.

### Microscopy for ASC speck formation

THP-1 differentiated macrophages were infected with anti-PstS1-coated ZsGreen expressing Mtb at MOI of 5 for 24hrs. For ASC staining, samples were fixed for 2hrs with 4% PFA at RT, then permeabilized in 2% BSA, 0.2% saponin, 0.1% TritonX100 for 15mins at RT. Samples were blocked in 2% BSA and incubated overnight at 4 C with anti-ASC/TSM1 (D2W8U), Rabbit mAb (1:200), from Cell Signaling (#67824). The samples were washed again and stained with Hoeschst (1:1000) and Alexa Fluor 647 conjugated goat anti-rabbit secondary antibody (Thermo Fisher; 1:300) for 1 h at RT. Images were obtained in one plane, using Evident Fluoview FV3000RS confocal microscope fitted with a 20x objective. Experiments were performed with duplicate wells per condition and repeated with at least 5 independent experiments. ASC specks per nuclei as determined by Hoeschst were quantified using Leica Imaris version10.1 analysis software following application of Gaussian filter.

### Mouse infection

Mice were purchased from the Jackson laboratories and bred at the UCSF animal facility. All procedures were performed in compliance with University of California San Francisco IACUC approved protocols. For pharmacological inhibition of NLRP3 inflammasome in vivo, C57BL/6 6-8 weeks old female and male mice were injected intraperitoneally with 10 mg/kg MCC950 (Invivogen) for 17 days starting at day −3 prior to aerosol infection. One day prior to Mtb infection mice were injected with PBS, Isotype control or p4-36 intraperitoneally. Mice were infected with Mtb strain HN878 by aerosol infection using a Glas-Col inhalation exposure chamber targeting an inoculum of ∼10-20 CFU. Lungs were harvested at day 14 post-infection; Homogenates of lung single cell suspension were plated on 7H11 agar and CFU was quantified after 21 days.

C57BL/6 and NLRP3 KO (B6.129S6-*Nlrp3*^tm1Bhk^/J – Jackson) of similar age and adjusted for sex were injected intraperitoneally with either PBS, Isotype control, p4-36 or p4-163 one day prior to Mtb infection. Mice were then aerosol challenged with ∼10-20 CFU Mtb strain HN878 using a Glas-Col inhalation exposure chamber as before. Lungs were harvested at day 14 post-infection for CFU quantification as before.

### Experiments with plasma from non-human primates

All plasma samples were provided by PAD and RAS from the Vaccine Research Center of NIAID, NIH. Samples were derived from unvaccinated (UNV), intradermally- (ID) and intravenously- (IV) vaccinated Rhesus macaques. Vaccine was with BCG strain Danish (see **Extended Data Table 1** for further details). Plasma from all groups were collected 4-5 weeks post-vaccination. Total IgG levels from plasma were measured using 1:1000 diluted plasma and a macaque total IgG ELISA kit (Abcam). Bacteria-specific immunoglobulin titers were measured by first coating ELISA plates overnight at 4C with the specified bacterial species (live *E. coli*-K12 or live BCG strain Pasteur, both OD=0.1, or heat-killed Mtb-HN878, total protein 2µg/mL). The following day, plates were washed, blocked with assay diluent and incubated with dilute plasma (1:25 for IgG1, 1:500 for IgM and 1:1000 for total IgG) at room temperature for 2hrs with shaking. After washing, plates were incubated with horseradish-peroxidase conjugated secondary antibodies against IgG or IgG1 (both Southern Biotech) or IgM (Sigma) and developed following manufacturer’s instructions as above.

For induction of IL-1β from macrophages, plasma was diluted 1:1000 in cell-culture medium and incubated with Mtb-HN878 to form immune complexes as for mAbs as above. BMDMs from B6 mice were infected with Mtb/Ab immune complexes (MOI 5) for 1hr at 37C and the assay otherwise performed as described above.

For passive transfer to mice experiments, age- and sex-matched mice were injected with 0.5mL undiluted macaque plasma via the intra-peritoneal route 24hrs prior to aerosol infection with Mtb-HN878 as described above. Plasma from each donor macaque was injected into one of 2-4 mice. At day 14 post-infection, mice were euthanized and lung homogenates plated for CFU as above.

### Statistical analyses

Graphpad Prism 10.4.2 was used for statistical analysis. Comparisons of two groups used one-way ANOVA followed by Dunnett’s test Comparisons of 3 or more groups use one-way or 2-way ANOVA and correction for multiple comparisons unless otherwise specified in the Figure legends.

## Acknowledgements

We thank Joel Ernst (UCSF) for sharing BCG Pasteur strain, Mtb-H37Rv, Mtb-HN878 and Mtb-ZsGreen. Mtb strains were confirmed to express PDIM. We thank Ernst lab members for helpful advice on experimental procedures, and the UCSF Core immunology Lab for help with flow cytometry. We thank Arun Prakash and Sara Suliman (both UCSF) for initial *Nlrp3* mice and primary human monocytes respectively. We thank Daniel Bachovchin (Memorial Sloan Cancer Center) for sharing THP-1 Caspase-1 KO cells. We thank members of the Javid lab for technical support for experiments and animal husbandry.

## Author contributions

RB and BJ designed experiments and analyzed data. RB performed the majority of experiments with assistance from NS, AF, TD. Confocal microscopy was performed by NS. Monoclonal antibodies and mutants thereof were generated by AW, LA, KR, NTF, PAM. BCG-immunized sera from macaques were provided by PAD and RAS. BJ obtained funding for the work. RB and BJ wrote the manuscript with input from other authors.

## References

1. Mangtani, P. et al. Protection by BCG vaccine against tuberculosis: a systematic review of randomized controlled trials. Clin Infect Dis 58, 470–480 (2014).

2. Chen, T. et al. Capsular glycan recognition provides antibody-mediated immunity against tuberculosis. J Clin Invest 130, 1808–1822 (2020).

3. Irvine, E.B. et al. Fc-engineered antibodies promote neutrophil-dependent control of Mycobacterium tuberculosis. Nature Microbiology 9, 2369–2382 (2024).

4. Li, H. et al. Latently and uninfected healthcare workers exposed to TB make protective antibodies against Mycobacterium tuberculosis. Proc Natl Acad Sci U S A 114, 5023–5028 (2017).

5. Liu, Y., et al. Features and protective efficacy of human monoclonal antibodies targeting Mycobacterium tuberculosis arabinomannan. JCI Insight (2023).

6. Lu, L.L. et al. A Functional Role for Antibodies in Tuberculosis. Cell 167, 433–443 e414 (2016).

7. Watson, A. et al. Human antibodies targeting a Mycobacterium transporter protein mediate protection against tuberculosis. Nature communications 12, 602 (2021).

8. Zimmermann, N. et al. Human isotype-dependent inhibitory antibody responses against Mycobacterium tuberculosis. EMBO Mol Med 8, 1325–1339 (2016).

9. Tait, D.R. et al. Final Analysis of a Trial of M72/AS01E Vaccine to Prevent Tuberculosis. New England Journal of Medicine 381, 2429–2439 (2019).

10. Li, H. & Javid, B. Antibodies and tuberculosis: finally coming of age? Nature reviews. Immunology (2018).

11. Schroder, K. & Tschopp, J. The inflammasomes. Cell 140, 821–832 (2010).

12. Beckwith, K.S. et al. Plasma membrane damage causes NLRP3 activation and pyroptosis during Mycobacterium tuberculosis infection. Nature communications 11, 2270 (2020).

13. Dorhoi, A. et al. Activation of the NLRP3 inflammasome by Mycobacterium tuberculosis is uncoupled from susceptibility to active tuberculosis. Eur J Immunol 42, 374–384 (2012).

14. Wong, K.W. & Jacobs, W.R., Jr. Critical role for NLRP3 in necrotic death triggered by Mycobacterium tuberculosis. Cellular microbiology 13, 1371–1384 (2011).

15. Deets, K.A. & Vance, R.E. Inflammasomes and adaptive immune responses. Nat Immunol 22, 412–422 (2021).

16. Koo, I.C. et al. ESX-1-dependent cytolysis in lysosome secretion and inflammasome activation during mycobacterial infection. Cellular microbiology 10, 1866–1878 (2008).

17. Master, S.S. et al. Mycobacterium tuberculosis prevents inflammasome activation. Cell host & microbe 3, 224–232 (2008).

18. Mishra, B.B. et al. Mycobacterium tuberculosis protein ESAT-6 is a potent activator of the NLRP3/ASC inflammasome. Cellular microbiology 12, 1046–1063 (2010).

19. Rastogi, S. & Briken, V. Interaction of Mycobacteria With Host Cell Inflammasomes. Frontiers in Immunology 13 (2022).

20. Rastogi, S., Ellinwood, S., Augenstreich, J., Mayer-Barber, K.D. & Briken, V. Mycobacterium tuberculosis inhibits the NLRP3 inflammasome activation via its phosphokinase PknF. PLoS pathogens 17, e1009712 (2021).

21. Sousa, J. et al. Mycobacterium tuberculosis associated with severe tuberculosis evades cytosolic surveillance systems and modulates IL-1beta production. Nature communications 11, 1949 (2020).

22. Taxman, D.J., Huang, M.T. & Ting, J.P. Inflammasome inhibition as a pathogenic stealth mechanism. Cell host & microbe 8, 7–11 (2010).

23. Mayer-Barber, K.D. et al. Caspase-1 independent IL-1beta production is critical for host resistance to mycobacterium tuberculosis and does not require TLR signaling in vivo. Journal of immunology 184, 3326–3330 (2010).

24. Rathinam, V.A. & Fitzgerald, K.A. Inflammasome Complexes: Emerging Mechanisms and Effector Functions. Cell 165, 792–800 (2016).

25. Krishnananthasivam, S. et al. An anti-LpqH human monoclonal antibody from an asymptomatic individual mediates protection against Mycobacterium tuberculosis. NPJ Vaccines 8, 127 (2023).

26. He, Y., Hara, H. & Nunez, G. Mechanism and Regulation of NLRP3 Inflammasome Activation. Trends in biochemical sciences 41, 1012–1021 (2016).

27. Coll, R.C. et al. A small-molecule inhibitor of the NLRP3 inflammasome for the treatment of inflammatory diseases. Nature medicine 21, 248–255 (2015).

28. Lu, L.L., Suscovich, T.J., Fortune, S.M. & Alter, G. Beyond binding: antibody effector functions in infectious diseases. Nature reviews. Immunology 18, 46–61 (2018).

29. Dahan, R. et al. Therapeutic Activity of Agonistic, Human Anti-CD40 Monoclonal Antibodies Requires Selective FcgammaR Engagement. Cancer Cell 29, 820–831 (2016).

30. Liu, S.D. et al. Afucosylated antibodies increase activation of FcγRIIIa-dependent signaling components to intensify processes promoting ADCC. Cancer immunology research 3, 173–183 (2015).

31. Yamin, R. et al. Fc-engineered antibody therapeutics with improved anti-SARS-CoV-2 efficacy. Nature 599, 465–470 (2021).

32. Smith, P., DiLillo, D.J., Bournazos, S., Li, F. & Ravetch, J.V. Mouse model recapitulating human Fcγ receptor structural and functional diversity. Proceedings of the National Academy of Sciences 109, 6181–6186 (2012).

33. Darrah, P.A. et al. Prevention of tuberculosis in macaques after intravenous BCG immunization. Nature 577, 95–102 (2020).

34. Darrah, P.A. et al. Airway T cells are a correlate of i.v. Bacille Calmette-Guerin-mediated protection against tuberculosis in rhesus macaques. Cell host & microbe 31, 962–977 e968 (2023).

35. Irvine, E.B. et al. Humoral correlates of protection against Mycobacterium tuberculosis following intravenous BCG vaccination in rhesus macaques. iScience 27, 111128 (2024).

36. Junqueira, C. et al. FcgammaR-mediated SARS-CoV-2 infection of monocytes activates inflammation. Nature 606, 576–584 (2022).

37. Maglione, P.J., Xu, J., Casadevall, A. & Chan, J. Fc gamma receptors regulate immune activation and susceptibility during Mycobacterium tuberculosis infection. Journal of immunology 180, 3329–3338 (2008).

38. Grace, P.S. et al. Antibody-Fab and -Fc features promote Mycobacterium tuberculosis restriction. Immunity (2025).

39. McEwan, W.A. et al. Intracellular antibody-bound pathogens stimulate immune signaling via the Fc receptor TRIM21. Nat Immunol 14, 327–336 (2013).

40. Ball, D.P. et al. Caspase-1 interdomain linker cleavage is required for pyroptosis. Life Sci Alliance 3 (2020).

41. Nieto-Caballero, V.E. et al. History of tuberculosis disease is associated with genetic regulatory variation in Peruvians. PLoS Genetics 20, e1011313 (2024).

42. Prados-Rosales, R. et al. The Type of Growth Medium Affects the Presence of a Mycobacterial Capsule and Is Associated With Differences in Protective Efficacy of BCG Vaccination Against Mycobacterium tuberculosis. The Journal of infectious diseases 214, 426–437 (2016).

