## Supplementary Data for "Early Antibody-Mediated Immunity to Tuberculosis in Mice Requires NLRP3"

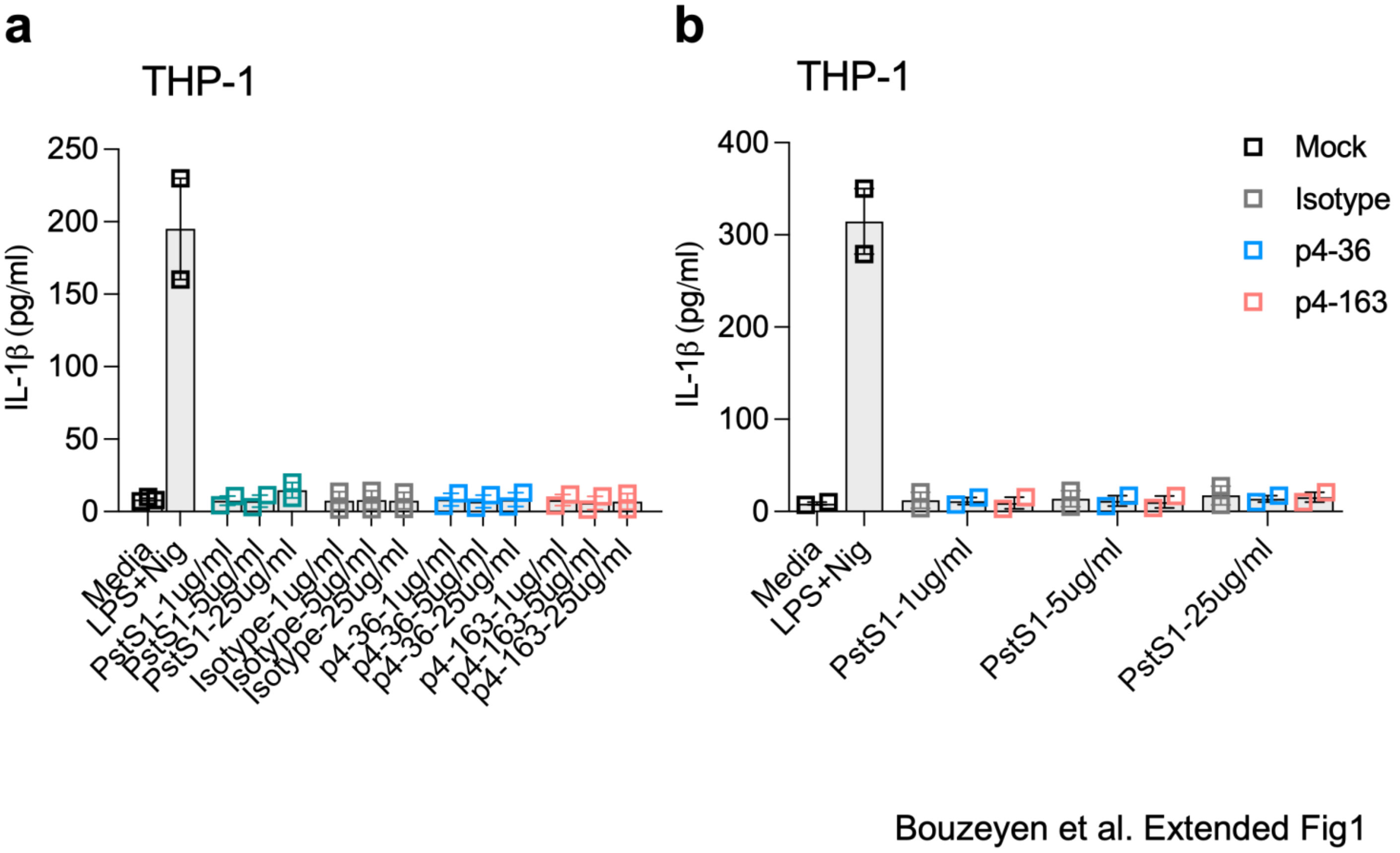


**Extended Data Fig. 1. PstS1 or PstS1-specific mAbs alone or as immune complexes are insufficient to stimulate IL-1β from macrophages.** Left: PMA-differentiated THP-1 macrophages were stimulated with PstS1 alone or PstS1-specific monoclonal antibodies p4-36 or p4-163 (or IgG1 isotype control) alone at indicated concentrations. Culture supernatant was collected after 24hrs and IL-1β measured by ELISA. LPS+ Nigericin used as a positive control for inflammasome activation. Right: immune complexes of PstS1 at indicated concentrations and PstS1-specific mAbs (25µg/mL) were formed for 1hr at 37C and then used to stimulate THP-1 macrophages as in first panel. IL-1β measured from culture supernatant after 24hrs as before.

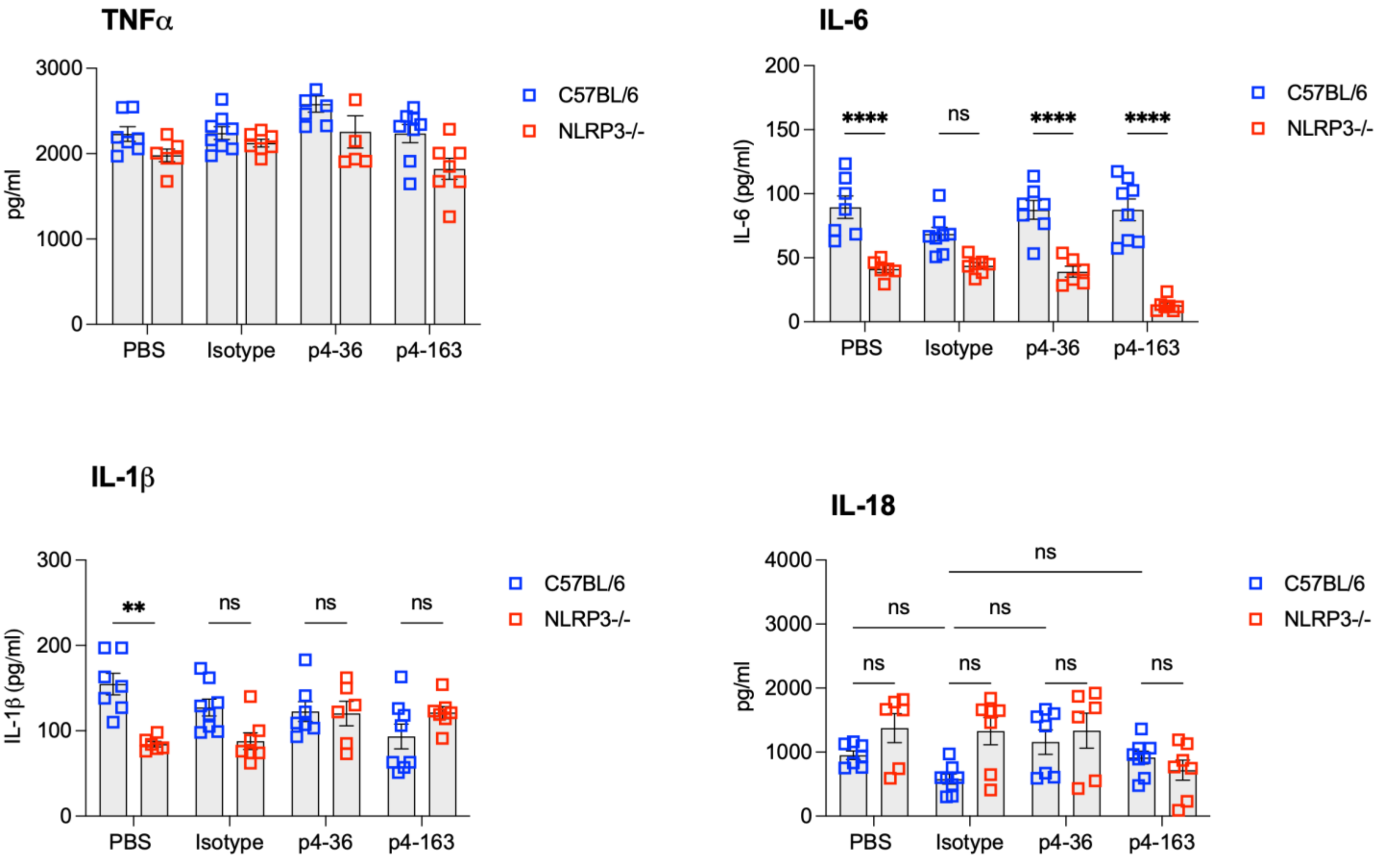


Bouzeyen et al. Extended Fig 2

**Extended Data Fig. 2. Cytokine analysis from lung homogenates of Mtb-infected mice.** Data refers to experiment in Fig. 4e,f. WT C57BL/6 or *Nlrp3*-deleted mice were injected with PstS1-specific mAbs, isotype control or PBS one day prior to aerosol infection with Mtb-HN878. At d14, mice were euthanized and lung homogenates plated for CFU as shown in Fig. 4. Homogenates were also tested for indicated cytokines by ELISA. Bars represent means +/- SEM. Each dot represents data from a single animal. **p<0.01, ****p<0.0001, nsp>0.05 by two-way ANOVA followed by post-hoc correction for multiple comparisons.

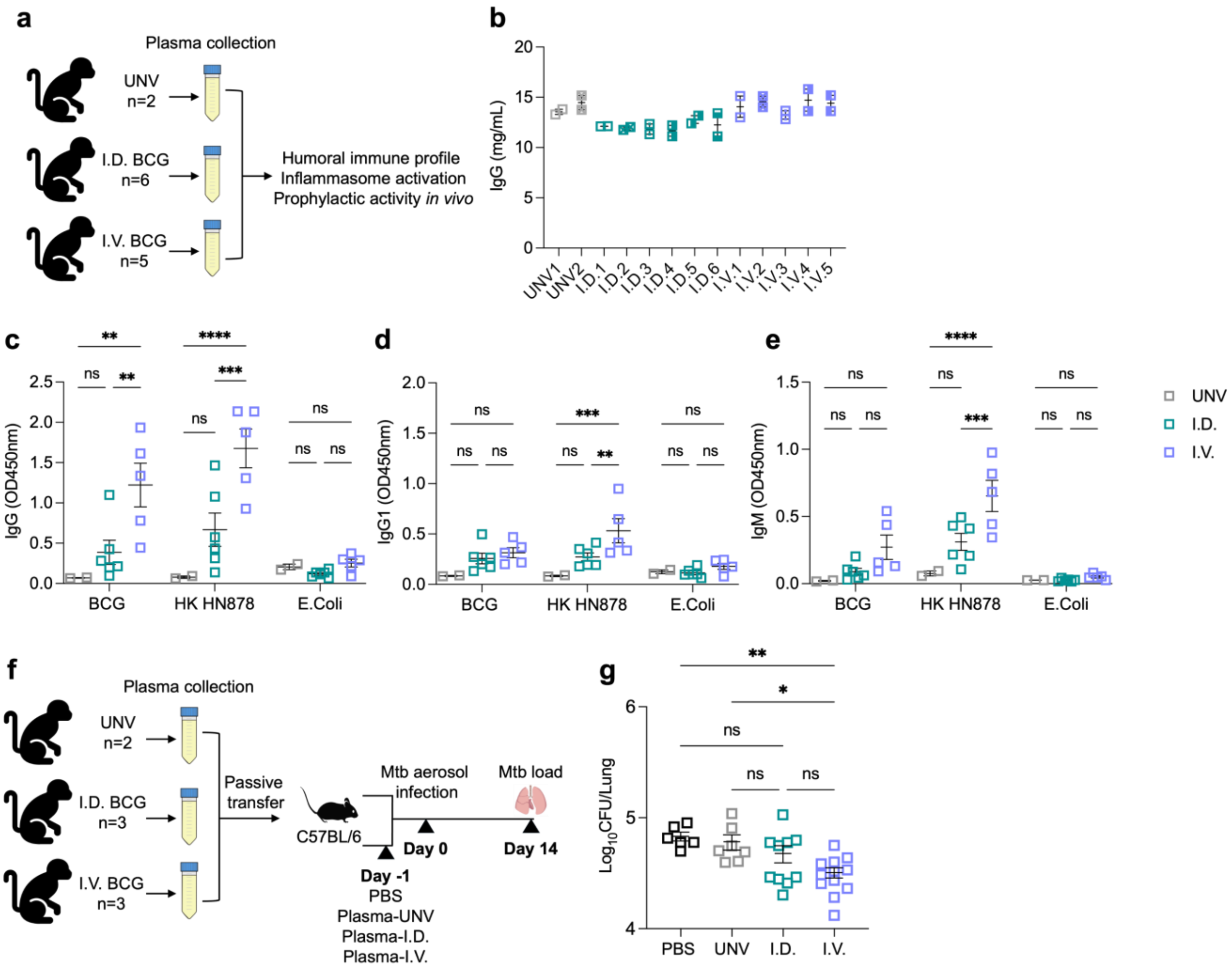
a
Plasma collection
UNV n=2
I.D. BCG n=6
I.V. BCG n=5
Humoral immune profile
Inflammasome activation
Prophylactic activity in vivo
b
IgG (mg/mL)
UNV1 UNV2 I.D.1 I.D.2 I.D.3 I.D.4 I.D.5 I.D.6 I.V.1 I.V.2 I.V.3 I.V.4 I.V.5
c
IgG (OD450nm)
d
IgG1 (OD450nm)
e
IgM (OD450nm)
BCG
HK HN878
E.Coli
UNV
I.D.
I.V.
f
Plasma collection
UNV n=2
I.D. BCG n=3
I.V. BCG n=3
Passive transfer
C57BL/6
Mtb aerosol infection
Mtb load
Day -1
Day 0
Day 14
PBS
Plasma-UNV
Plasma-I.D.
Plasma-I.V.
g
$Log_{10}$CFU/Lung
PBS UNV I.D. I.V.

**Extended Data Fig. 3. Passive transfer of plasma from IV-BCG immunized macaques protects mice from Mtb at day 14.** (a) Cartoon representing source of plasma from unvaccinated (UNV), intradermally BCG-immunized (ID) and intravenously BCG-immunized (IV) Rhesus macaques. (b) Total IgG in plasma from each macaque as measured by ELISA. IgG (c), IgG1 (d) and IgM (e) titers against live BCG, heat-killed Mtb-HN878 (HK HN878) and live *E. coli*-K12 (*E. coli*) from plasma samples. (f) Cartoon outling passive transfer experiment: plasma (0.5mL) from number of indicated macaques was injected into 2-4 C57/BL6 mice each 1 day prior to aerosol infection with Mtb-HN878 (~10 cfu/animal). At day 14 post-infection, mice were euthanized and lung homogenates plated for Mtb (g). Horizontal bars represent mean +/- SEM. Each dot represents data from one measurement (biological repliacate for ELISA or animal for cfu). *p<0.05, **p<0.01, ***p<0.001, ****p<0.0001, nsp>0.05 by two-way ANOVA with Šídák's post hoc test.

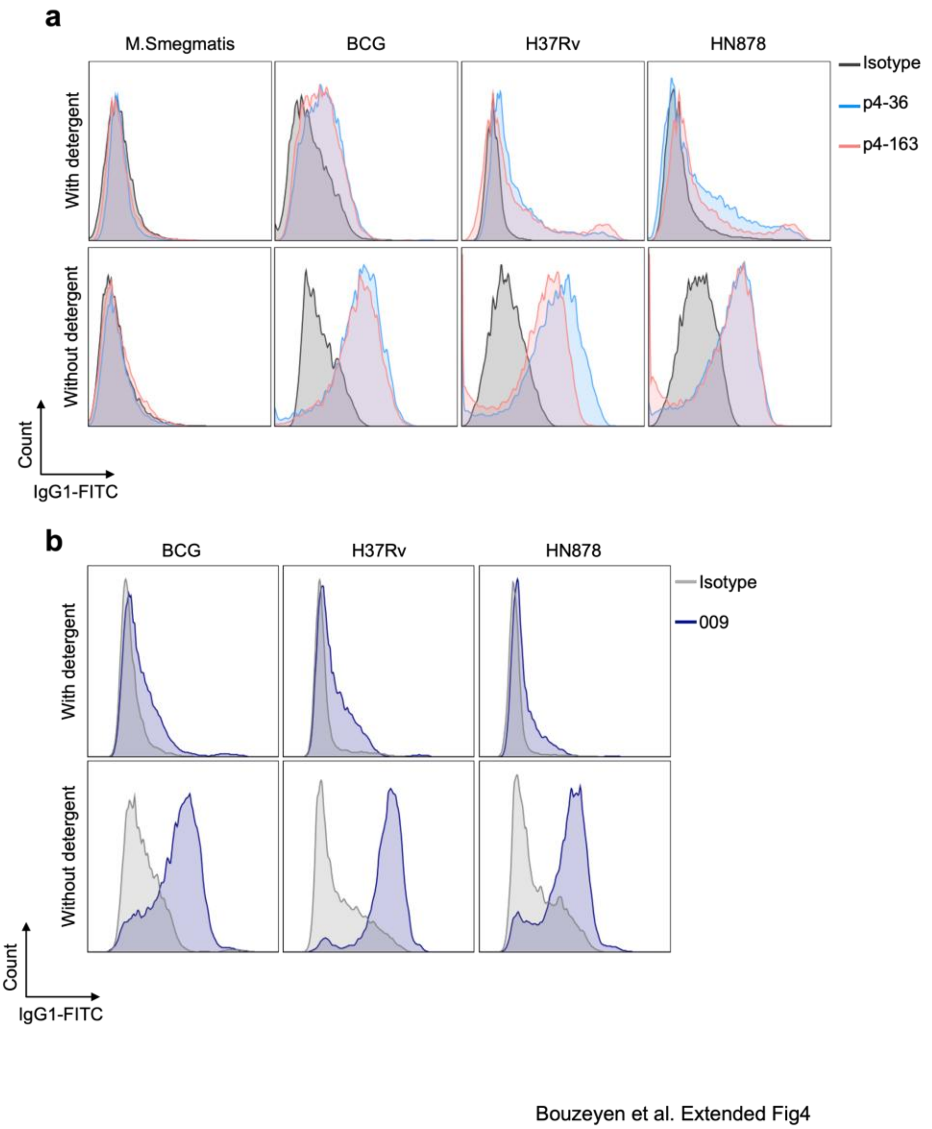


**Extended Data Fig. 4. Mtb-specific monoclonal antibody binding to live mycobacteria.** (a) Binding of PstS1-specific mAbs p4-36 and p4-163 or (b) LpqH-specific mAb 009 to live *M. smegmatis*, *M. bovis*-BCG (BCG), Mtb-H37Rv and Mtb-HN878. Bacteria were grown either with (upper panel) or without (lower panel) Tween detergent. Bacteria were suspended as single-cells then stained overnight with mAbs or isotype control (all 10µg/mL). Bacteria were then washed, stained with secondary antibody (mouse anti-human IgG1-FITC) and then fixed with 4% PFA before analysis by flow cytometry. Representative histograms from 3 biological replicates, performed at least twice are shown.

| NHP ID | Species | Origin | Sex | Date of birth | Vaccination Date | BCG Stock | Vaccination Route | Vaccination Administration | Vaccine volume | Vaccination Dose |
|---|---|---|---|---|---|---|---|---|---|---|
| DK05 (I.V.1) | Macaca mulatta | CPRC | M | 210611 | 240827 | 21.BCG.3.25.5.ws | Intravenous | Right leg, saphenous vein | 2ml (butterfly) | 3.69E+07 |
| CE8 (I.V.2) | Macaca mulatta | CPRC | M | 200517 | 240827 | 21.BCG.3.25.5.ws | Intravenous | Right leg, saphenous vein | 2ml (butterfly) | 3.69E+07 |
| CA7 (I.V.3) | Macaca mulatta | CPRC | M | 200609 | 240827 | 21.BCG.3.25.5.ws | Intravenous | Right leg, saphenous vein | 2ml (butterfly) | 3.69E+07 |
| BY2 (I.V.4) | Macaca mulatta | CPRC | M | 200811 | 240827 | 21.BCG.3.25.5.ws | Intravenous | Right leg, saphenous vein | 2ml (butterfly) | 3.69E+07 |
| CF9 (I.V.5) | Macaca mulatta | CPRC | M | 200614 | 240827 | 21.BCG.3.25.5.ws | Intravenous | Right leg, saphenous vein | 2ml (butterfly) | 3.69E+07 |
| DK06 (I.D.1) | Macaca mulatta | CPRC | M | 210802 | 240827 | 21.BCG.3.25.5.ws | Intradermal | Right and left upper arm | 200ul per arm | 3.69E+07 |
| CM1 (I.D.2) | Macaca mulatta | CPRC | M | 200717 | 240827 | 21.BCG.3.25.5.ws | Intradermal | Right and left upper arm | 200ul per arm | 3.69E+07 |
| BY4 (I.D.3) | Macaca mulatta | CPRC | M | 200819 | 240827 | 21.BCG.3.25.5.ws | Intradermal | Right and left upper arm | 200ul per arm | 3.69E+07 |
| CG1 (I.D.4) | Macaca mulatta | CPRC | M | 200714 | 240827 | 21.BCG.3.25.5.ws | Intradermal | Right and left upper arm | 200ul per arm | 3.69E+07 |
| BZ2 (I.D.5) | Macaca mulatta | CPRC | M | 200811 | 240827 | 21.BCG.3.25.5.ws | Intradermal | Right and left upper arm | 200ul per arm | 3.69E+07 |
| CK3 (I.D.6) | Macaca mulatta | CPRC | M | 200607 | 240827 | 21.BCG.3.25.5.ws | Intradermal | Right and left upper arm | 200ul per arm | 3.69E+07 |
| 08C102 (UNV 1) | Macaca mulatta | Covance | M | 080918 | Unvaccinated control | | | | | |
| 08C035 (UNV 2) | Macaca mulatta | Covance | M | 080501 | Unvaccinated control | | | | | |

**Extended Data Table 1. Data with regards to Rhesus macaques from which plasma was sourced for experiments.**
